# Genome mining of amylases and amylase inhibitors from *Streptomyces*

**DOI:** 10.64898/2026.01.13.699332

**Authors:** Andreas C Lawaetz, Alexa Gannon, Paul A Hoskisson

## Abstract

Most antibiotics are derived from *Streptomyces* bacteria and produced industrially via fermentations relying on food-grade feedstocks, which has a substantial environmental impact. Efficient utilisation of starch-rich organic wastes such as bread and potato waste as alternative feedstocks remains limited by incomplete knowledge of starch metabolism in *Streptomyces.* Here, a genus-wide analysis of 295 *Streptomyces* strains was performed, identifying 645 amylases grouped into pullulanases, α-1,4-amylases, and cyclomaltodextrin-like amylases, alongside biosynthetic gene clusters encoding amylase inhibitors such as acarbose and tendamistat. Structural and sequence analyses revealed that inhibitor resistance arises from subtle modifications in the α-amylase catalytic pocket. This comprehensive analysis of amylases and inhibitors provides a foundation for engineering inhibitor-resistant enzymes tailored to diverse starch substrates, facilitating the development of sustainable, starch-based *Streptomyces* fermentations.

**Impact statement:** Pharmaceutical manufacturing contributes substantially to global carbon emissions, with feedstocks used in natural product fermentations being a major factor. Replacing conventional feedstocks with organic waste streams, such as starch-rich bread or potato waste, could reduce the environmental impact, but two key challenges remain: production strains may lack the enzymes required to efficiently degrade alternative substrates, and nutrient changes may trigger carbon catabolite repression, reducing product yields. While previous studies have linked carbohydrate-active enzymes with biosynthetic gene clusters, no comprehensive analysis has focused specifically on starch degradation in *Streptomyces*. To fill this gap, a genus-wide analysis of amylases and amylase inhibitors across 295 *Streptomyces* genomes was carried out. This created an atlas of 645 amylases which may serve as the foundation for the targeted selection of enzymes optimised for catabolising organic waste from various sources, thereby improving the sustainably of industrial natural product production such as antibiotics.

**Data summary:** Scripts used along with supplementary files and figures can be accessed through GitHub at https://github.com/ALawaetz/Amylases_and_amylase_inhibitors_in_Streptomycetes.

## Introduction

The majority of antibiotics and other bioactive natural products are generated by complex biosynthetic pathways found naturally in *Streptomyces* bacteria and are produced industrially in large scale fermentations (Hoskisson and Seipke, 2020). Pharmaceutical manufacturing has a substantial impact on the environment and accounts for 5% of the UK’s carbon footprint with a large contribution coming from the carbon intensive processed sugars and lipids currently used as feedstocks in antibiotic production systems (Lenzen et al., 2020). One strategy to enhance the sustainability of pharmaceutical manufacturing is to replace current feedstocks with organic waste products. Globally, ∼1.3 billion tons of food waste is produced each year representing a cheap and sustainable energy source that could become part of a circular bioeconomy instead of ending up in landfills (Fritsch et al., 2017; Kumar et al., 2023). Bread waste, for example, is comprised of 50-70% starch which could potentially be fed into antibiotic fermentations (Tomasik and Horton, 2012; Kumar et al., 2023).

Changing feedstocks for bioactive natural products, such as antibiotics in production strains, however, is not straightforward as primary metabolism is intimately linked to biosynthesis, which is in turn subject to carbon catabolite repression (Fernández-Martínez and Hoskisson, 2019). To rationally engineer cells to metabolise desired carbon sources while maintaining high antibiotic titres, a thorough understanding of metabolic networks is required. Hence, substantial efforts have gone into correlating the repertoire of biosynthetic gene clusters (BGCs) with that of carbohydrate active enzymes (CAZymes) in many *Streptomyces* genomes (Vela Gurovic et al., 2021; Otani, Udwary and Mouncey, 2022; Ríos-Fernández, Caicedo-Montoya and Ríos-Estepa, 2024). While CAZyme analyses use resources such as the Carbohydrate Active Enzymes database (Drula et al., 2022) and annotation tools like dbCAN (Zheng et al., 2023a) and provide valuable insights into the metabolic breadth of organisms, they are limited by the level of functional detail they can offer. Many CAZyme categories are broad and encompass proteins with diverse enzymatic activities. For example, the glycoside hydrolase category 13 (GH13) family contains enzymes with activities as diverse as amylases, sucrases, and trehalose synthases. Whilst efforts are underway to further subdivide CAZyme families into more homogeneous, functionally coherent groups (Drula et al., 2022), these refinements remain incomplete. Therefore, to bioinformatically assess the capacity of streptomycetes to metabolise starch specifically, further characterisation beyond standard CAZyme annotation is required.

Starch is a naturally occurring polymer and many bacteria, including species of *Streptomyces,* encode and secrete amylases, which enable them to break down and catabolise starch (Tomasik and Horton, 2012). Starch consists of long chains of glucose residues covalently linked through α-1,4-glycosidic bonds and may further be branched through α-1,6 linkages (amylose and amylopectin, respectively; Tomasik and Horton, 2012). In the natural microenvironments of streptomycetes, other starch-like polysaccharides may be encountered including pullulan (composed of maltotriose units linked by α-1,6 bonds) and cyclomaltodextrins (cyclic oligomers of α-1,4-linked glucose residues). Concordantly, *Streptomyces* genomes have been annotated as encoding amylases, pullulanases, and cyclomaltodextrinases (Tomasik and Horton, 2012). Starch is among the most abundant and energy dense polymers in the environment (Tomasik and Horton, 2012) and fierce competition between microbes to access this resource is likely. Unsurprisingly, species of *Streptomyces* synthesise amylase inhibitors including carbohydrate-based compounds like acarbose, acarviostatin, and trestatin and small peptide inhibitors like tendamistat, Gaim, and Scaim (Aschauer et al., 1983; Rehm et al., 2009; Rockser and Wehmeier, 2009; Guo et al., 2012; Tanoeyadi et al., 2023).

In this study, combining bioinformatic resources including dbCAN (Zheng et al., 2023b), SignalP (Teufel et al., 2022), hmmer (Eddy, 2011), and GATOR (Cediel-Becerra et al., 2025) was used to assess the distribution of amylases and amylase inhibitors across 295 *Streptomyces* genomes, providing a broad representation of the starch catabolic diversity of the genus. It was found that pullulanases and α-1,4-acting amylases are almost ubiquitously present, whereas only a subset of strains encode a third, cyclomaltodextrin-like amylase. The total number of amylases per genome ranged from zero to six, and there was a positive correlation between strains encoding higher numbers of amylases and those encoding amylase inhibitors. BGCs encoding amylase inhibitors were typically found alongside multiple amylases, including variants with amino acid motifs known to provide resistance to inhibition (Tanoeyadi et al., 2023).

## Results

To assess the prevalence of amylases in *Streptomyces* species, dbCAN (Zheng et al., 2023a) was used to search 295 genomes, broadly encompassing the genetic diversity of the genus (Kiepas, 2025), for proteins belonging to CAZyme family GH13 (Drula et al., 2022). Results were subsequently filtered for protein secretion signals using SignalP (Teufel et al., 2022) to yield 645 amylases which formed three discrete phylogenetical clades (Figure 1A). Analysis of conserved domains found that the three clades constituted pullulanases (clade 1), α-1,4-amylases (clade 2), and cyclomaltodextrin-like amylases (clade 3; Figure 1B). To assess the distribution of each type of amylase across the *Streptomyces* genus, the presence of each was mapped to each genome using a phylogenetic tree displaying the evolutionary relationship between all 295 genomes (Figure 2). It was found that most strains encoded pullulanases and α-1,4-amylases (87% and 89%, respectively) whereas only 22% of strains had a cyclomaltodextrin-like amylase in their genome. The total number of amylases per genome varied between zero and six.

**Figure 1:**
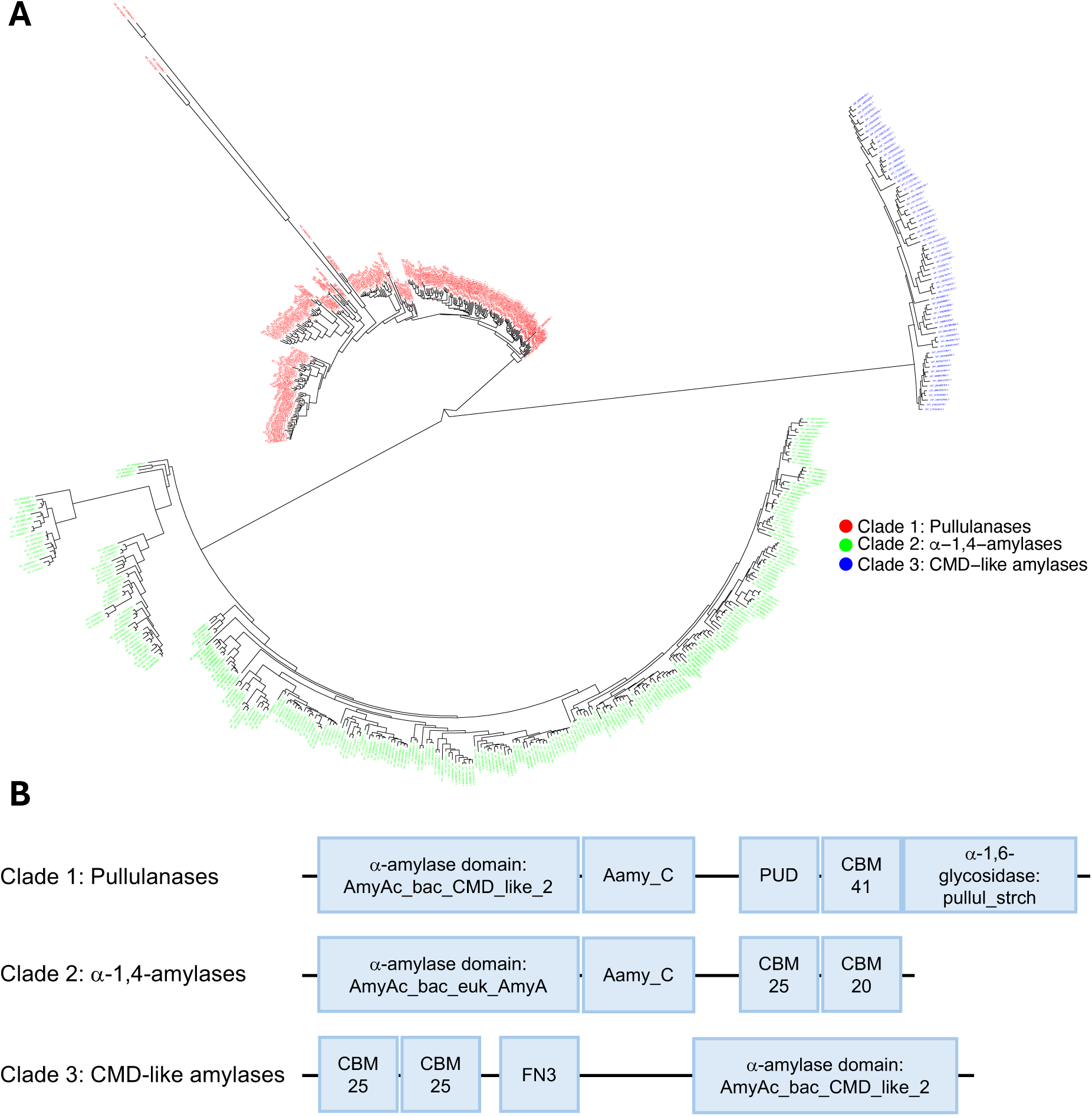
Streptomycetes possess three types of amylases. A total of 645 amylases were found across 295 *Streptomyces* genomes. Phylogenetic analysis showed that amylases in streptomycetes cluster into three major clades (A). Conserved domain analysis of representative proteins from each clade found that clade 1 consisted of pullulanases, clade 2 of α-1,4-acting amylases, and clade 3 of cyclomaltodextrin (CMD)-like amylases (B). AmyAc_bac_CMD_like_2: α-amylase catalytic domain found in bacterial cyclomaltodextrinases; Aamy_C: maltogenic C-terminal amylase domain; PUD: pullulanase associated domain; CBM: carbohydrate binding module; pullul_strch: pullulanase type α-1,6-glycosidase domain; AmyAc_bac_euk_AmyA: α-1,4-amylase catalytic domain found in bacterial and eukaryotic α-amylases; FN3: Fibronectin type 3 domain. Domains not drawn to scale.

**Figure 2:**
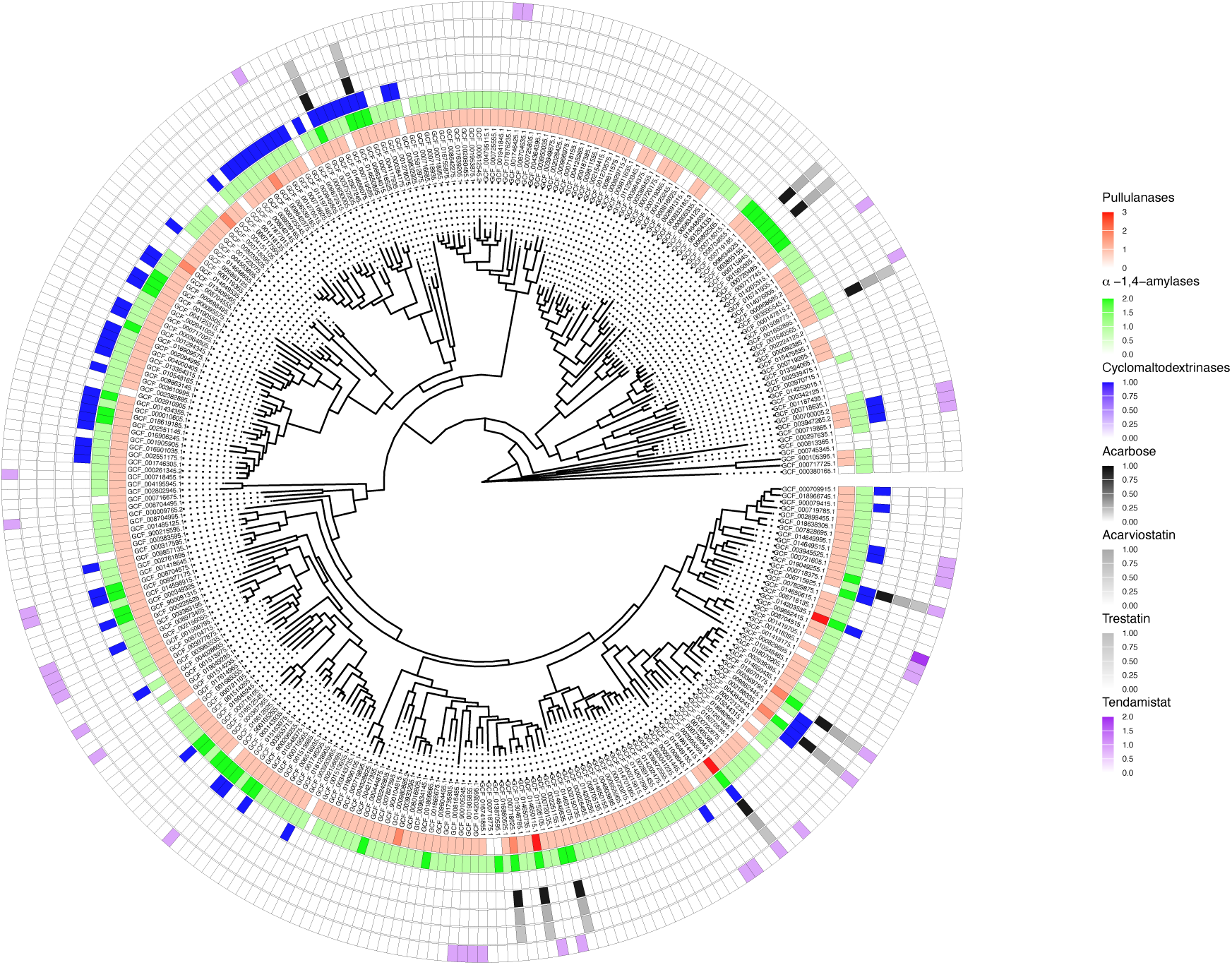
Amylases frequently co-occur with amylase inhibitors in *Streptomyces* genomes. The evolutionary relationship between 295 *Streptomyces* species were inferred from whole genome phylogeny and the prevalence of amylases and amylase inhibitors is visualised using a colour scale (white indicates absence; darker colours indicate higher prevalence). Annotations shown in the legend (right panel), listed from top to bottom, correspond to the concentric circles around the phylogenetic tree from the innermost to the outermost. Pullulanases (innermost, red) and α-1,4-amylases (green) are found almost ubiquitously throughout the *Streptomyces* genus, whereas only a subset of species encode cyclomaltodextrin-like amylases (blue). Genomes with higher numbers of amylases frequently coincide with those encoding amylase inhibitors, including acarbose (black), acarviostatin (darkgrey), trestatin (grey), and tendamistat (purple).

Previous studies have reported BGCs encoding amylase inhibitory molecules acarbose, acarviostatin, and trestatin located adjacent to amylase gene arrays (Rockser and Wehmeier, 2009; Guo et al., 2012; Tanoeyadi et al., 2023). To test if strains encoding amylase inhibitors generally possess more amylase genes, a hmmsearch (Eddy, 2011) with a tendamistat HMM profile was used to analyse the genomes and found 38 species encoding tendamistat homologs. Using GATOR (Cediel-Becerra et al., 2025), 12 species were identified having BGCs encoding acarbose, acarviostatin, or trestatin (Figure 2 and 3). These clusters were typically located adjacent to multiple amylase genes (numbers ranging from one to four; Figure 3). Acarbose, acarviostatin, and trestatin are all pseudooligosaccharides but they vary in the number of C_7_N-aminocyclitol units they contain (Tanoeyadi et al., 2023). It was found that all acarbose-like gene clusters harboured a common set of genes required for C_7_N-aminocyclitol synthesis (see materials and methods) but differed in the domain architecture of a pseudoglycosyltransferase responsible for the non-glycosidic C-N bonding in pseudooligosaccharides (Tsunoda et al., 2022). Further variation was seen in their neighbouring genes including homologs of the acarbose importer AcbFGH (Wehmeier and Piepersberg, 2004; Brunkhorst et al., 2005) which was absent in some clusters (Figure 3). A Mann-Whitney U test revealed a strong positive correlation between genomes with higher numbers of amylases and those with acarbose BGCs (r = 0.78; p = 4.95x10^-7^; Figure 4A). A similar analysis found a significant, yet weaker correlation between higher number of amylases and genomes encoding tendamistat (r = 0.27; p = 2.93x10^-3^) (Figure 4B). A Chi-square test further revealed a positive but weak correlation between cyclomaltodextrin-like amylases and tendamistat (phi = 0.16; p = 0.009).

**Figure 3:**
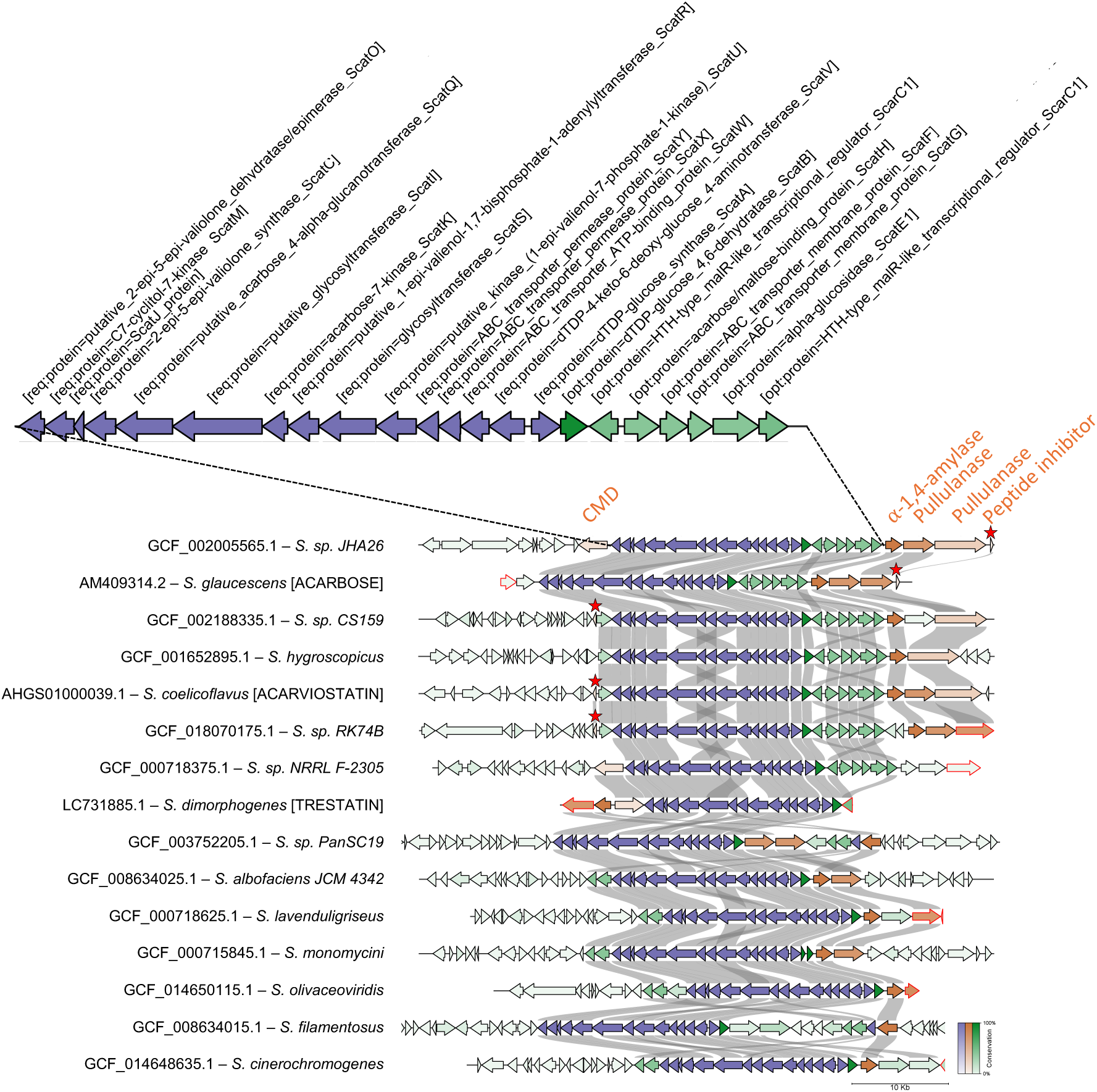
Acarbose-like BGCs possess between one and four amylases in their gene-neighbourhoods. GATOR identified 15 genomes (12 from the representative list of 295 genomes plus the acarviostatin, acarbose, and trestatin BGCs already described in literature). Core genes required for the synthesis of aminocyclitol are shown in blue, amylases are shown in orange, and other genes in green. Tendamistat-like peptide inhibitors are shown in orange and marked with a red star. Gene level conservation between clusters are shown by colour intensity (more intense colour for higher level of conservation). Genes with red edges indicate contig edges. CMD: cyclomaltodextrinase.

**Figure 4:**
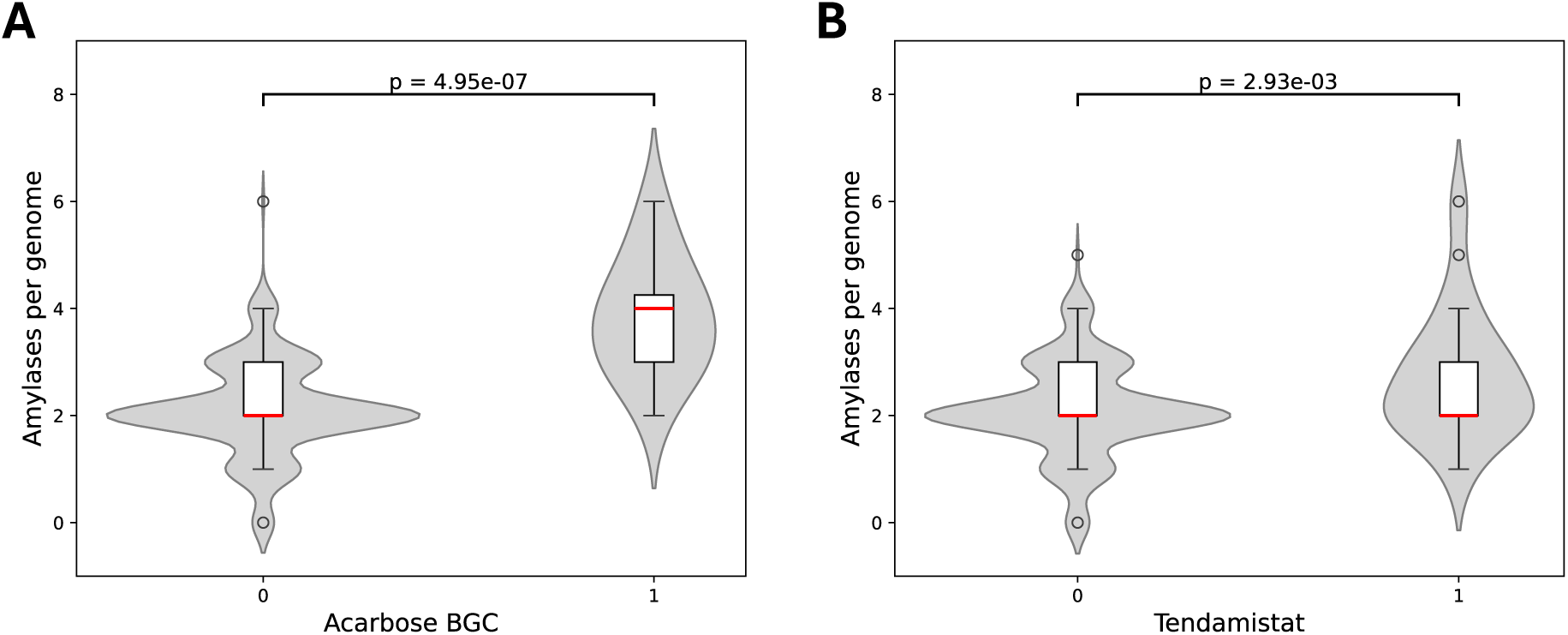
*Streptomyces* species encoding amylase inhibitors possess more amylase genes. Violin-boxplots showing the distribution of amylases per genome in strains with and without acarbose BGCs (A) or tendamistat (B).

Previous studies in *S. glaucescens* and *S. dimorphogenes* (Tanoeyadi et al., 2023) pairing *in silico* analysis with mutational screening identified a catalytic pocket in α-amylases where substitution of a histidine with alanine or asparagine was sufficient to confer resistance to acarbose (Tanoeyadi et al., 2023). To evaluate whether this might be a general feature of amylases associated with acarbose-like BGCs, a multiple sequence alignment (MSA) was constructed (Supplementary file S5) and found seven amylases across six genomes with acarbose-like BGCs having a histidine to asparagine substitution at this catalytic site. Although the substitution also occurred in 11 genomes lacking such BGCs (Supplementary Figure S5), statistical analysis found it 30-fold enriched in species harbouring acarbose-like clusters (p = 1.93e-07). Interestingly, not all species encoding acarbose had an amylase with the described histidine to arginine substitution (Supplementary Figure S1), suggesting that acarbose resistance may be achieved through other mechanisms.

To find possible acarbose-resistant amylases, a MSA was constructed for each clade of amylases (pullulanases, α-1,4-amylases, and cyclomaltodextrin-like amylases), identifying numerous amino acid motifs significantly enriched in species harbouring acarbose-like BGCs (Supplementary Files S2-S4). In acarbose BGC associated α-1,4-amylases it was found that they possess several amino acid changes including a H/Y to G and a D to T substitution (Figure 5A). Using AlphaFold3 (Jumper et al., 2021) to model the structures of an acarbose-associated α-1,4-amylase (WP_077796428.1) and a non-acarbose-associated amylase from the same clade (WP_103530170.1), a noticeable difference in the architecture of the catalytic pocket (Matsuura et al., 1984; Buisson et al., 1987; Uitdehaag et al., 1999; Tanoeyadi et al., 2023) was found between the two groups (Figure 5B). Importantly, using CB-DOCK2 (Liu et al., 2022) to model the binding of acarbose, it was found that the inhibitor is predicted to bind both amylases but in different orientations (Figure 5B). To assess, *in silico,* the effect of the H235G and D293T substitutions in the acarbose-associated amylase, the amino acids were reverted to H and D, respectively, and it was found that this alone was enough to widen the catalytic pocket and change the binding configuration of acarbose similar to that predicted for the non-acarbose-associated amylase (Figure 5B). A similar conversion was observed by making the reciprocal substitution in the non-acarbose-associated amylase (Supplementary Figure S2). Importantly, modelling of an α-1,4-amylase (WP_150242730) from an acarbose BGC without a pullulanase having the H to N acarbose-resistance-substitution also displayed an inhibitor-resistant acarbose binding phenotype that could be reversed through a single G238H substitution (Supplementary Figure S5). It was hypothesised that reorienting the binding of acarbose within the catalytic pocket might be a general way of acquiring acarbose resistance and therefore the binding of acarbose to an acarbose-associated pullulanase (WP_064455463) was modelled. It was found that acarbose was predicted to bind in a similar way (Supplementary Figure S3) to that determined for the putatively acarbose-resistant α-1,4-amylase, with the imino moiety of acarbose oriented away from the region of the binding pocket known to be instrumental in establishing acarbose resistance (Tanoeyadi et al., 2023). The asparagine to histidine conversion appears to result in the reversion of this group of pullulanases from an inhibitor resistant to an inhibitor sensitive conformation (Tanoeyadi et al., 2023) and the modelling predicts a reorientation of acarbose binding (Supplementary Figure S3). A lysine residue directly adjacent to this site is conserved in both pullulanases and α-1,4-amylases (Supplementary File S5), and as described previously, the orientation of this residue’s side chain varies considerably between resistant and sensitive amylases (Tanoeyadi et al., 2023). Similar observations were found in α-1,4-amylases (Supplementary Figure S4), likely attributable to altered hydrogen bonding networks in the catalytic pocket (Figure 5).

**Figure 5:**
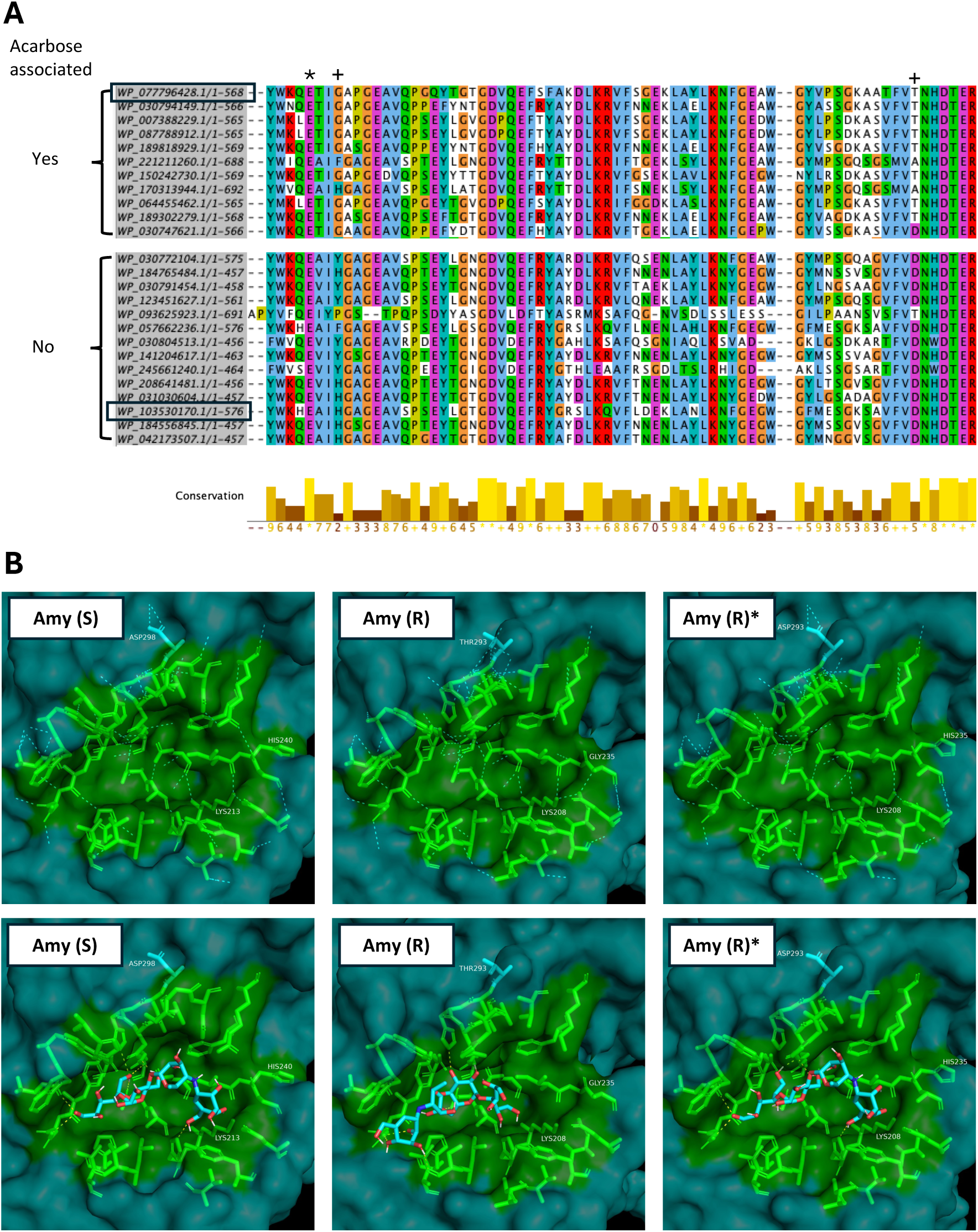
Predicted binding sites of acarbose differs between amylases that are located in acarbose BGCs and amylases that are not. MSA shows an enrichment of GLY235 and THR293 (+) in alpha-1,4-amylases located in acarbose BGCs. Both residues are found within the conserved catalytic pocket close to the catalytic residue glutamate (*) (A). The catalytic pockets (residues highlighted in green) are shown for a putatively acarbose-sensitive amylase [Amy (S) – accession: WP_103530170] found in a genome without acarbose and a putatively resistant amylase [Amy (R) – accession: WP_077796428] found within an acarbose BGC [proteins marked by black squares in (A)]. Amy (R)* shows the structure of Amy (R) where GLY235 and THR293 have been converted to HIS and ASP, identical to the residues found at these positions in Amy (S). Hydrogen bonds between residues within the catalytic pocket are shown with dashed cyan lines and hydrogen bonds between acarbose and the proteins are shown with dashed yellow lines (bottom panel). LYS213/208, previously described as involved in inferring acarbose resistance is highlighted (B).

The acarbose and acarviostatin BGCs of *S. glaucescens* and *S. coelicoflavus* encode in addition to pseudooligosaccharides also small proteinacous amylase inhibitors, Gaim and Scaim (Guo et al., 2012; Tanoeyadi et al., 2023). These peptides contain the conserved amylase inhibitor domain (smart00783) also found in tendamistat. Tendamistat is known to inhibit eukaryotic amylases (Vertesy et al., 1984; Wiegand, Epp and Huber, 1995), however, little is known about its activity against prokaryotic amylases. GATOR identified three additional *Streptomyces* species that harbour acarbose BGCs encoding small peptides with amylase inhibitor domains (Figure 3). To assess potential inhibition of *Streptomyces* amylases, interactions between Gaim and three pullulanases from *Streptomyces sp. JHA26* (GCF_002005565.1), two enoded within the acarbose BGC and one outside the cluster, were modelled using ClusPro (Kozakov et al., 2017). Gaim was predicted to bind in the α-amylase catalytic pocket of the non-BGC-associated pullulanase as well as one of the two BGC-associated pullulanases, resembling the resolved interaction between tendamistat and porcine pancreatic amylase (Figure 6; (Wiegand, Epp and Huber, 1995). No comparable binding was predicted for the second BGC-associated pullulanase, likely due to protruding side chains within its binding pocket that obstruct Gaim binding (Figure 6).

**Figure 6:**
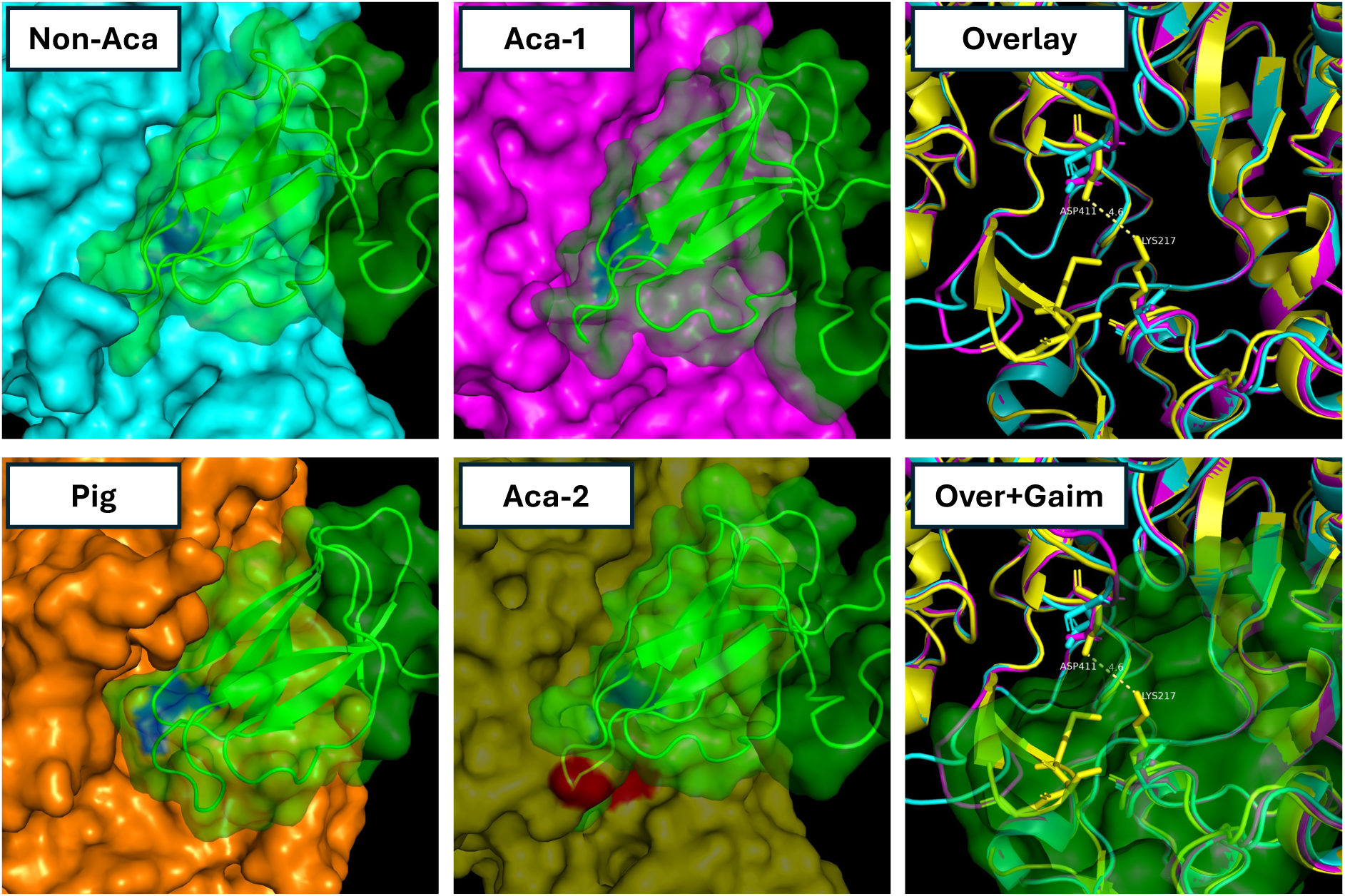
Structure predictions and protein-protein docking analyses suggest obstructed binding of proteinaceous amylase inhibitor Gaim to an acarbose-associated pullulanase from *Streptomyces sp. JHA26.* Gaim is predicted to bind two pullulanases from *Streptomyces sp. JHA26*: one encoded within the acarbose BGC (Aca-1 [magenta]; WP_077796430.1) and one located outside the BGC (Non-Aca [cyan]; WP_077796080.1). Gaim (green) encoded within the acarbose BGC is predicted to bind in the starch binding pocket containing the catalytic aspartate and glutamate residues (blue), in a manner similar to the binding of tendamistat (green) to porcine pancreatic amylase (orange; Pig). Alignment of the second acarbose BGC-encoded pullulanase (Aca-2 [transparent yellow]; WP_077796429.1) with the Gaim-bound non-acarbose-associated pullulanase reveals three protruding amino acid residues (red) that are predicted to obstruct Gaim binding. An overlay of the three *Streptomyces* amylases highlights these residues (Overlay), including weak electrostatic interactions between D411 and K217 which creates a closed pocket confirmation that sterically clashes with Gaim binding (Over+Gaim).

## Discussion

Streptomycetes are known to inhabit a wide range of ecological niches and exhibit extensive genome plasticity particularly in their repertoires of BGCs and CAZymes (Norte, Avitia-Dominguez and Rozen, 2025). In accordance, this work reveals substantial diversity in both the number and types of amylases across the genus. It was found that *Streptomyces* amylases broadly cluster into three major clades comprised of pullulanases, α-1,4-amylases and cyclomaltodextrin-like amylases. While all three clades are homologous in their α-amylase domains, they differ in their adjacent carbohydrate binding domains. This likely confers substrate specificity toward starches of varying composition, such as those derived from different plant species or fungal sources. It was also found that *Streptomyces* species frequently harbour BGCs encoding amylase inhibitors, such as acarbose and tendamistat, alongside inhibitor resistant amylases. It is known that acarbose not only inhibits amylases but may also function as a “carbophor” for the producing organism (Hemker et al., 2001; Wehmeier and Piepersberg, 2004). In this role, acarbose is covalently linked to glucose or dextrins through the acarviosyltransferase AcbD and then transported into the cytoplasm through the ABC transporter AcbFGH-MsiK (Wehmeier and Piepersberg, 2004; Brunkhorst et al., 2005). Notably, this work indicates that acarbose resistance is more likely achieved through reconfiguration of acarbose binding within the alpha-amylase catalytic pocket rather than exclusion of the inhibitor. This suggests that acarbose-resistant amylases may participate in the carbophor cycle, enabling acarbose-producing organisms to sequester oligosaccharides, that might otherwise be freely accessible to competing organisms, by covalently linking them to acarbose, thereby restricting uptake to organisms possessing the dedicated importer.

The atlas of amylases presented in this work could serve as a useful resource for the targeted selection of enzymes for heterologous overexpression in industrial antibiotic production strains. While further functional characterisation of individual amylase clades and subclades is required, this resource may facilitate the optimisation of sustainable antibiotic fermentations by deploying amylases with substrate specificities tailored to organic waste streams of different origins. Moreover, this approach may provide a useful source of bacterial strains that produce novel alpha-glucosidase inhibitors for therapeutic use in diabetes mellitus (Moelands et al., 2018).

In the context of engineering starch-fuelled industrial cell factories, it is important to consider the potential repertoire of endogenous amylase inhibitors encoded by a given production strain. Amylase inhibitors are frequently found in *Streptomyces* bacteria, and care must be taken to ensure that any host strain does not produce inhibitors that act against a heterologously expressed amylase, as this could compromise manufacturing efficiency. *Streptomyces hygroscopicus*, for example, is used industrially for the large-scale production of a range of bioactive compounds, including rapamycin (Park et al., 2010). Improving the sustainability of fermentations involving this organism using organic waste substrates, such as bread or potato waste, would be of considerable environmental value (Ribeiro and Silva, 2022; Kumar et al., 2023). However, as this analysis shows, *S. hygroscopicus* (GCF_001652895.1) encodes an acarbose BGC, and this may potentially interfere with amylase activity during fermentation. Strategies to overcome this challenge could be to either delete this cluster or, alternatively, overexpress an acarbose-resistant amylase. The latter approach may be more broadly applicable, as overexpressing genes often is easier than deleting them. Moreover, considering the ongoing antibiotic resistance crisis and the need to discover and industrially deploy new antibiotics, production pipelines should be readily adaptable to newly identified producer strains. Rather than repeatedly facing the challenging task of deleting gene clusters encoding amylase inhibitors in newly characterised strains, it may be more efficient to develop a small set of broadly applicable overexpression vectors. Furthermore, as this and previous work suggests, creating inhibitor-resistant amylases may be achieved through minimal sequence modifications, including single amino acid substitutions. It may hence be possible to develop a toolbox of inhibitor-resistant amylases with optimised substrate specificities toward starches from diverse sources, which could be readily integrated into a wide range of natural product-producing *Streptomyces* strains.

In summary, this work highlights the coordinated evolution of amylases and amylase inhibitors in *Streptomyces,* revealing how starch utilisation and inhibitor resistance are tightly integrated. These findings provide insight into how producing organisms balance nutrient acquisition with competitive exclusion and offer a framework for engineering inhibitor-resistant, starch utilising strains for sustainable antibiotic fermentations.

## Materials and methods

### Phylogenetic analysis of amylases in *Streptomyces*

A database of 295 *Streptomyces* genomes (Kiepas, 2025); accession numbers found in Supplementary File S6) were used for the analysis and the protein fasta files for each genome were downloaded with the NCBI Datasets command-line tools [script: Supplementary File S10]. Amylases were found by using dbcan (version: 5.1.2) (Zheng et al., 2023a) with the option CAZyme_annotation [script: Supplementary File S7]. Results were subsequently filtered for GH13 proteins where this was the recommended result (i.e. agreement between hmmer and DIAMOND). Proteins whose recommended result included multiple CAZyme groups, e.g. GH13 and/or CBM20 were also included in the analysis. Results that included subgroups, e.g. GH13_32 were included in the analysis simply as GH13 [script: Supplementary File S8]. SignalP (version: 6.0) (Teufel et al., 2022) was then used to select only proteins with secretion signals [script: Supplementary File S9], yielding a total of 645 amylases. A multiple sequence alignment of all amylases was carried out using Mafft (version: 7.525) (Madeira et al., 2024) [script: Supplementary File S11] and their phylogenetic relationship were determined using RAxML-NG (version: 1.2.2) (Kozlov et al., 2019) with 100 bootstrap replicates [script: Supplementary File S12]. A circular tree rooted at midpoint was drawn from the resulting output using R [script: Supplementary File S13]. Domain analysis was carried out using NLM’s conserved domain database (Wang et al., 2023) using representative proteins from each clade. Representative proteins were defined as the proteins most similar to the consensus sequence in each clade (script: Supplementary File S22). These were WP_062143504.1 (Clade 1), WP_218063397.1 (Clade 2), and WP_150491328.1 (Clade 3).

### Amylase inhibitors

Previously characterised BGCs encoding amylase inhibitors acarbose (Tanoeyadi et al., 2023), acarviostatin (Guo et al., 2012) and trestatin (Tanoeyadi et al., 2023) were used to search *Streptomyces* genomes using GATOR-GC (version: 0.9.0) (Cediel-Becerra et al., 2025) [script: Supplementary File S14]. GATOR-GC takes two lists of genes as input; a list of required genes and a list of optional genes. Required genes to search for acarbose BGC were the core genes required for aminocyclitol synthesis: gac[A, V, W, X, Y, U, S, R, K, I, Q, C, J, M, O]. The homologous genes in *S. coelicoflavus* ZG0656 and *S. dimorphogenes ATCC 31484* were used to search for acarviostatin and trestatin BGCs, respectively. A combined list of optional genes was used to search all clusters. This included all amylases and proteinaceous amylase inhibitors from the characterised BGCs in *S. glaucescens* (Rockser and Wehmeier, 2009; Tanoeyadi et al., 2023), *S. coelicoflavus* ZG0656 (Guo et al., 2012), and *S. dimorphogenes ATCC 31484* (Tanoeyadi et al., 2023), all 645 amylases identified in this study, and tendamistat from *S. tendae* (M28478.1). A genus-wide search for tendamistat-like peptides was carried out by building a hmmer profile (HMMER version: 3.4) from a tendamistat seed alignment (smart domain smart00783 from the NLM’s conserved domain database) and then using hmmsearch [script: Supplementary Files S15] with an E-value cutoff of 10^-10^ to find occurrences across the *Streptomyces* genus. Occurrences of amylases and amylase inhibitors were plotted using R around a circular tree rooted at midpoint displaying the evolutionary relationship between 295 *Streptomyces* genomes based on whole genome phylogeny (Kiepas et al), [script: Supplementary File S16].

### Multiple sequence alignment

Multiple sequence alignments of each identified clade of amylases were carried out with Mafft (version: 7.525) (Madeira et al., 2024) using the seed option (seed alignment from conserved α-amylase domain cd00551) [Supplementary File S11]. Aligned sequences were visualised with Jalview (version: 2.11.5.1) (Waterhouse et al., 2009).

### Statistical analysis

The distribution of the number of amylases per genome was tested using the scipy.stats shapiro package and found to not be normally distributed. A Mann-Whitney U test was hence applied to test whether the distributions of the number of amylases in genomes with and without acarbose BGC were significantly different from each other. All genomes carried either one or zero cyclomaltodextrin-like amylases and one or zero tendamistat genes and with two binary variables, a chi-square test was used to test the correlation between the two. To test for enrichment of aligned amino acids in amylases found in acarbose BGCs, a fisher exact test was applied and corrected for multiple testing using FDR (Benjamini-Hochberg) and Bonferroni. A p-value of 0.05 was used as cutoff for significance. All scripts used for statistical analysis can be found in Supplementary File S17 and S18.

### Protein structures and ligand binding modelling

AlphaFold 3 (Jumper et al., 2021) was used to model the 3-dimensional structure of proteins. CB-DOCK2 (Liu et al., 2022) and ClusPro 2.0 (Kozakov et al., 2017) were used to predict the binding of acarbose (PubChem CID 9811704) and Gaim to amylases, respectively. Default parameters were used in all cases. For simplicity, modelling of binding interactions between Gaim and pullulanases was done using the α-amylase domains only. As such, it cannot be excluded that amylase inhibitor resistance may be achieved through interdomain interactions obstructing Gaim binding. Open source PyMOL was installed using conda and used to visualise and annotate proteins and protein-ligand interactions [script: Supplementary files S19-S21]

## CRediT (Contributor roles taxonomy)

Conceptualization – ACL, PAH.

Data curation - ACL.

Formal analysis - ACL.

Funding acquisition – PAH.

Investigation - ACL.

Methodology – ACL, PAH.

Project administration – ACL, PAH.

Supervision - PAH.

Validation - ACL.

Visualization – ACL

Writing – original draft – ACL

Writing – review & editing – ACL, AG, PAH.

All authors gave final approval for publication and agreed to be held accountable for the work performed therein.

## Ethics

This work did not require ethical approval from a human subject or animal welfare committee.

## Data accessibility

The data relevant to the conclusions of this paper and supplementary material is deposited here https://github.com/ALawaetz/Amylases_and_amylase_inhibitors_in_Streptomycetes.

## Declaration of AI use

We have not used AI-assisted technologies in creating this article.

## Acknowledgements

We would like to thank Prof Iain S Hunter, Dr J T Munnoch, and Dr Steve Kendrew (GSK) for helpful discussions.

---

**Supplementary Figure S1:**
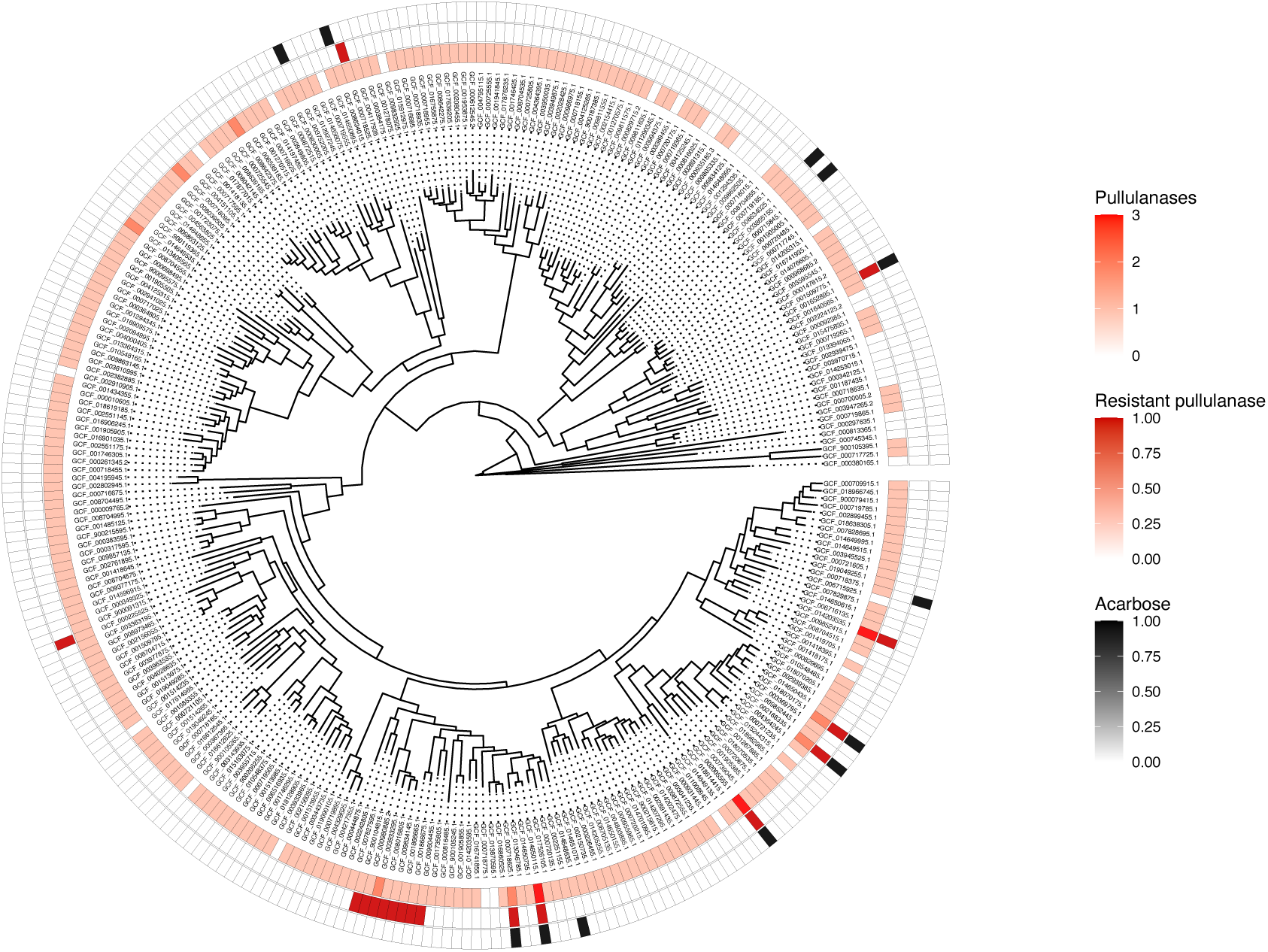
Amylase inhibitor resistant amylases are enriched in genomes encoding acarbose BGCs.

**Supplementary Figure S2:**
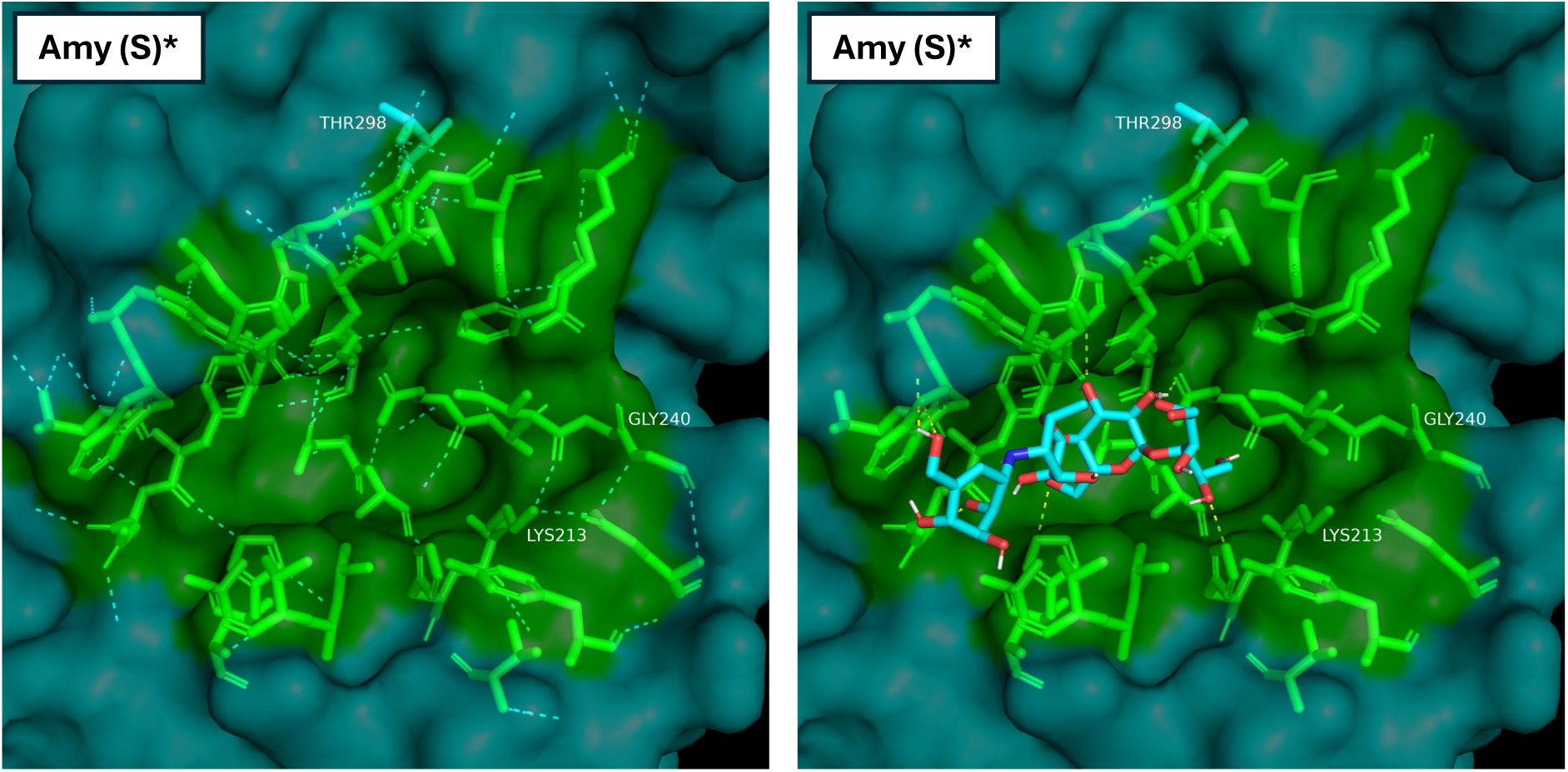
D298T and H240G changes the binding pattern of acarbose in a non-acarbose-associated amylase (WP_103530170) to that seen in acarbose-associated amylases.

**Supplementary Figure S3:**
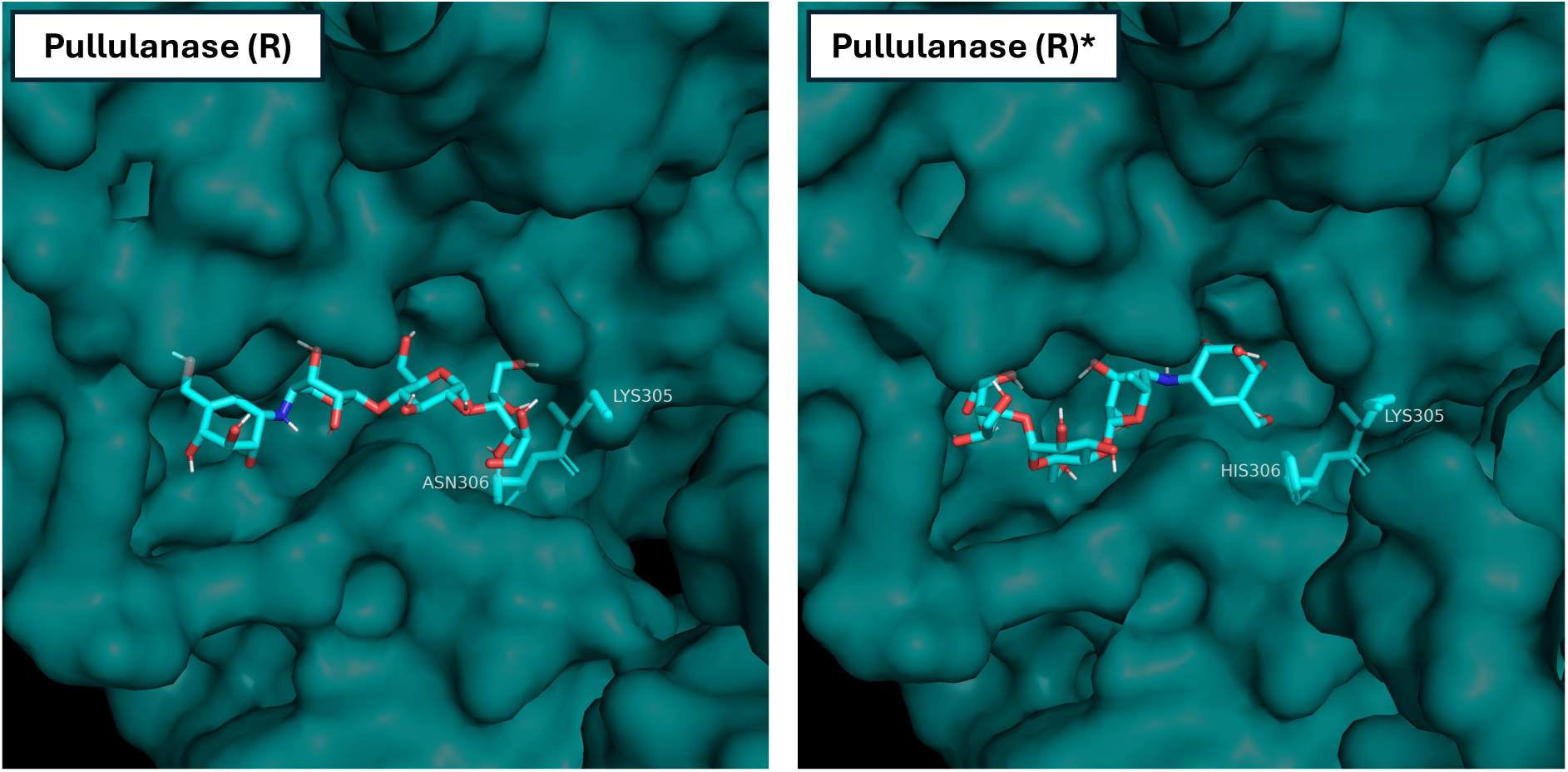
N306H reorients the predicted binding of acarbose in an inhibitor resistant pullulanase (WP_064455463) to a configuration similar to that seen in alpha-1,4-amylases found in genomes without acarbose.

**Supplementary Figure S4:**
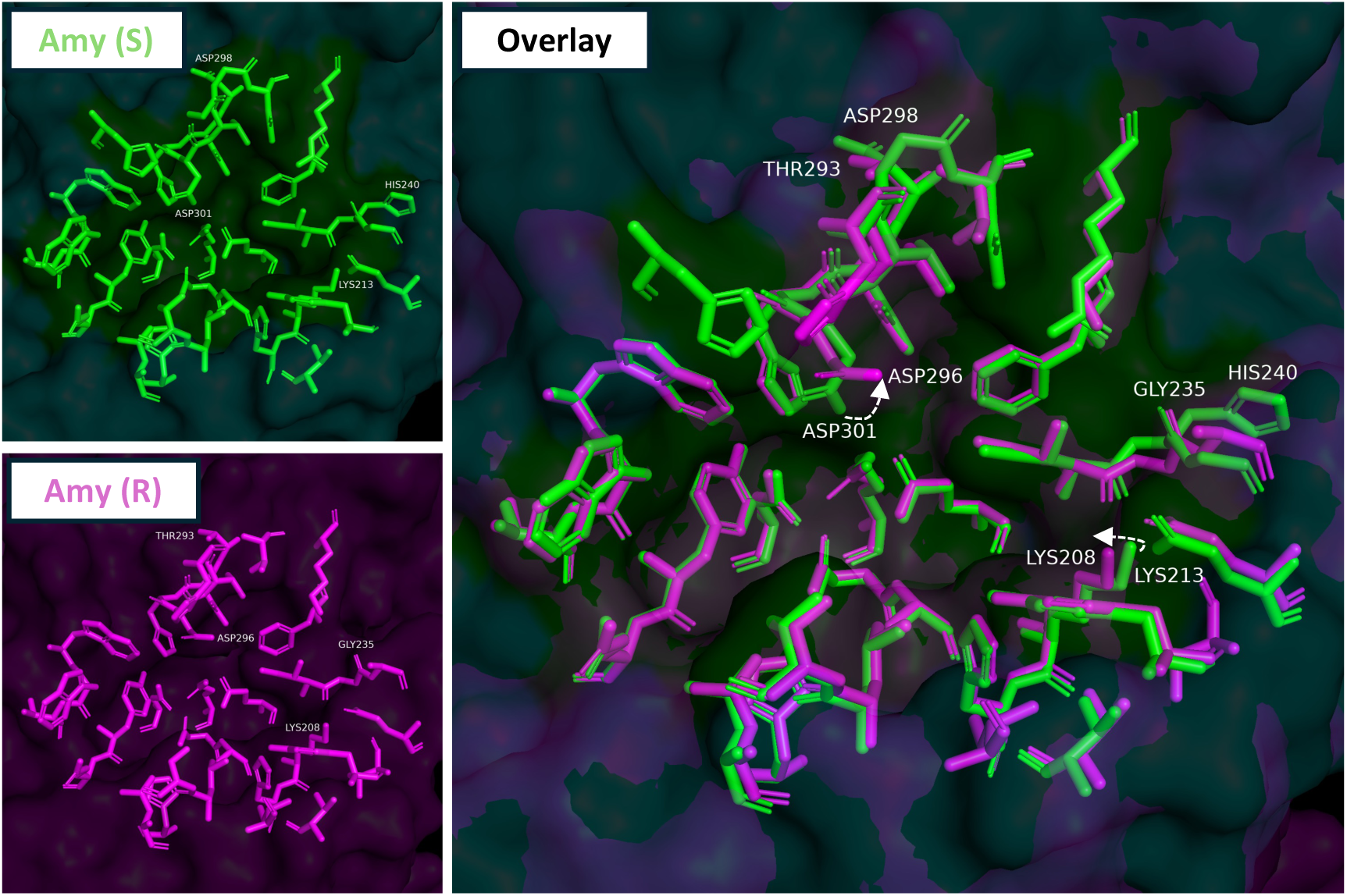
The architecture of the catalytic pocket differs between Amy (S) and Amy (R). Amy (S) [WP_103530170; green] and Amy (R) [WP_077796428; magenta] varies on residue 293/298 and 235/240 which might be causative of the reorientation of D296/301 and K208/213 resulting in the narrower binding pocket found in Amy (R).

**Supplementary Figure S5:**
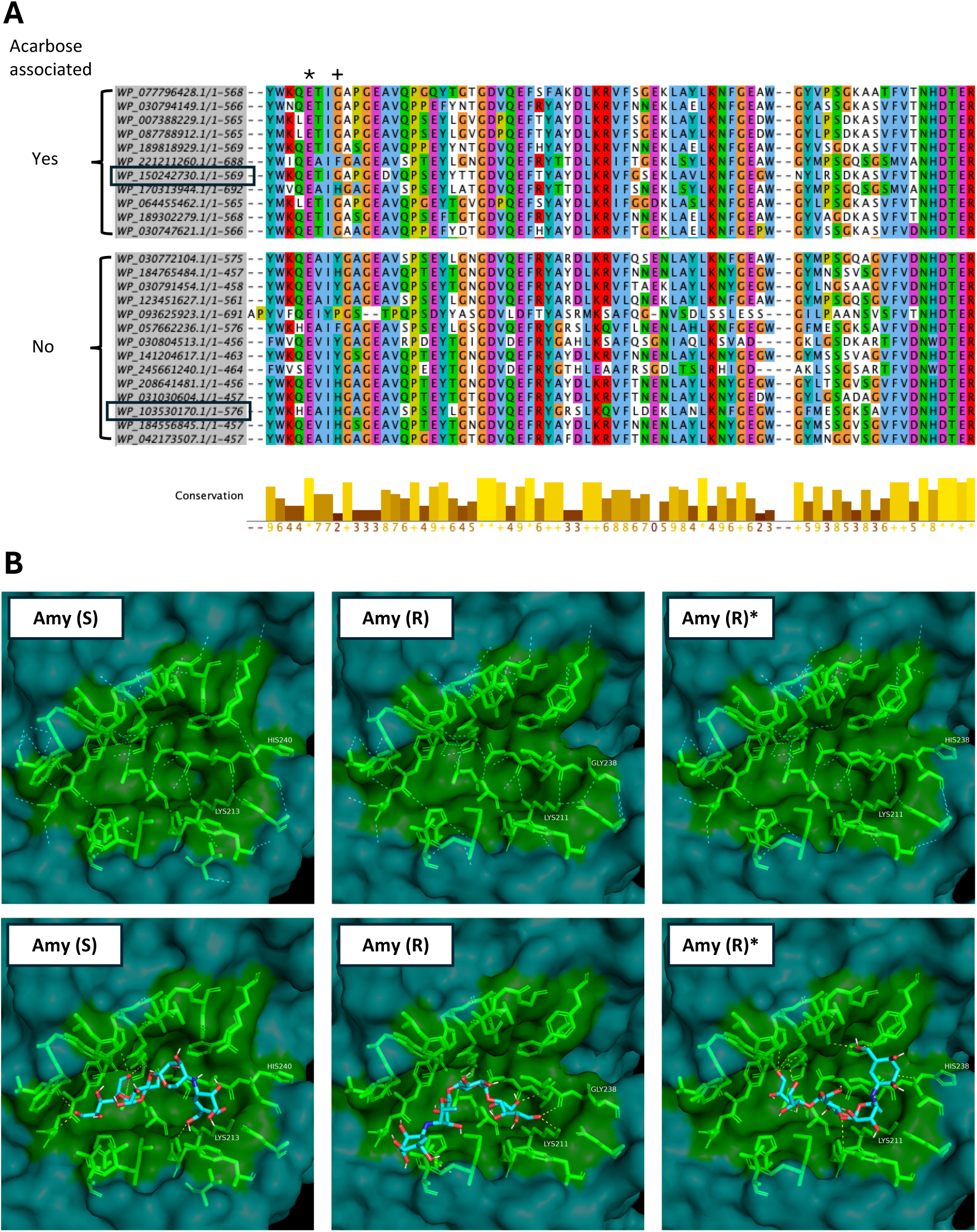
G238H reorients acarbose binding in an α-1,4-amylase from an acarbose BGC without an acarbose resistant pullulanase. MSA shows an enrichment of GLY238 (+) in alpha-1,4-amylases located in acarbose BGCs (A). The catalytic pockets (residues highlighted in green) are shown for a putatively acarbose-sensitive amylase [Amy (S) – accession: WP_103530170] found in a genome without acarbose and a putatively resistant amylase [Amy (R) – accession: WP_150242730] found within an acarbose BGC [proteins marked by black squares in (A)]. Amy (R)* shows the structure of Amy (R) where GLY238 have been converted to HIS, identical to the residue found at this positions in Amy (S). Hydrogen bonds between residues within the catalytic pocket are shown with dashed cyan lines and hydrogen bonds between acarbose and the proteins are shown with dashed yellow lines (bottom panel). LYS213/211, previously described as involved in inferring acarbose resistance is highlighted (B).

